# Regulation of co-translational mRNA decay by PAP and DXO1 in Arabidopsis

**DOI:** 10.1101/2024.12.20.629663

**Authors:** Marie-Christine Carpentier, Anne-Elodie Receveur, Adrien Cadoudal, Rémy Merret

**Author notes:** Present address: Rémy Merret, Institut de Biologie Moléculaire des Plantes, UPR2357 CNRS, Université de Strasbourg, 12, rue du Général Zimmer, 67084 Strasbourg, France.

## Abstract

**Background:** mRNA decay is central in the regulation of mRNA homeostasis in the cell. The recent discovery of a co-translational mRNA decay pathway (also called CTRD) has changed our understanding of the mRNA decay process. This pathway has emerged as an evolutionarily conversed mechanism essential for specific physiological processes in eukaryotes, especially in plants. In Arabidopsis, this pathway is targeted mainly by the exoribonuclease XRN4. However, the details of the molecular regulation of this pathway are still unclear.

**Results:** In this study, we first tested the role of the 3ʹ-phosphoadenosine 5ʹ-phosphate (PAP), an inhibitor of exoribonucleases in the regulation of CTRD. Using 5’Pseq approach, we discovered that FRY1 inactivation impaired XRN4-CTRD activity. Based on this finding, we demonstrated that exogenous PAP treatment stabilizes CTRD mRNA targets. Furthermore, we also tested the implication of the exoribonuclease DXO1 in CTRD regulation. We found that DXO1, another exoribonuclease sensitive to PAP, is also involved in the CTRD pathway, especially by targeting NAD^+^-capped mRNAs. DXO1 specifically targets mRNAs linked to stress response.

**Conclusions:** Our study provides further insights into the regulation of CTRD in Arabidopsis and demonstrates that other exoribonucleases can be implicated in this pathway.

## BACKGROUND

The tight regulation of messenger RNA (mRNA) abundance is necessary to ensure correct development and stress response in eukaryotic organisms. Changes in mRNA abundance can be induced not only by modulating the transcription level but also by modulating the mRNA decay rate. In eukaryotes, mRNA decay can occur from both extremities of the transcript. The 3’ to 5’ mRNA decay is targeted by the RNA exosome complex or SOV (suppressor of varicose) in Arabidopsis (1,2). The 5’ to 3’ mRNA decay takes place after the deadenylation and decapping processes. The uncapped mRNA is then targeted by an exoribonuclease (XRN1 in *Saccharomyces cerevisiae*, XRN4 in *Arabidopsis thaliana*) (3,4).

For many years, it was proposed that mRNA decay only occurs after the last round of translation. However, several studies have suggested that translated mRNAs can also be targeted for degradation (5). The discovery of the 5’-3’ co-translational mRNA decay (also called CTRD) validated this hypothesis. Pioneering evidence was demonstrated in yeast where uncapped mRNAs were purified from polysomal fractions (6). This work was then followed by several other studies with the development of Next-Generation Sequencing (NGS) strategies based on the capture of 5’ monophosphate decay intermediates (also called 5’Pseq) (7–12). These approaches revealed that mRNA decay intermediates follow a three-nucleotide periodicity. This periodicity is explained by the tracking of the last translating ribosome by the exoribonuclease. The mRNA decay rate is determined by the ribosome translocation rate codon per codon. Thus, it was proposed that in addition to mRNA decay analysis, these 5’Pseq approaches can also reveal ribosome dynamics at nucleotide resolution (8,11–17). The common feature identified in all 5’Pseq datasets is a 5’P reads accumulation 16 to 17 nucleotides upstream of the stop codons, explained by the slow translation termination step (11). The dynamic of this accumulation is also used as a proxy of CTRD activity (10,18–25).

This pathway appears to be evolutionarily conserved as it has been described in many eukaryotic and prokaryotic organisms such as bacteria, yeast, human and plants (11,16,19,21,22,24–26). In addition, this pathway was demonstrated to be essential for several physiological contexts. In mammalian cells, CTRD is involved in the regulation of tubulin mRNA to maintain the abundance of soluble tubulin in the cell (27–29). In yeast, this pathway has been shown to be important for the osmotic stress response (30). In plants, this pathway is present in at least 10 angiosperms with conserved features (19) and is regulated throughout Arabidopsis seedling development (21). During heat stress, CTRD is also activated and has been proposed to be essential for thermotolerance (31,32). In tomato, CTRD allows the regulation of circadian rhythm transcripts (22). Recently, the contribution of this pathway in the general mRNA turnover was also determined in Arabidopsis, revealing that this pathway is a major determinant in mRNA turnover in both shoot and root (20). Factors involved in the regulation of CTRD have also been described in plants. The RNA-binding protein LARP1 (La- Related protein 1) is involved in the targeting of XRN4 to polysomes especially during heat stress condition (31). CBP80, a large subunit of the cap binding complex is also involved in the regulation of CTRD (24). More recently, Pelota and Hbs1, two translation-related ribosome rescue factors, were proposed to be suppressors of CTRD in Arabidopsis (18). However, details about the molecular mechanisms that trigger CTRD are still rare in the literature.

XRN family members were first described in yeast and were found to have two members: ScXRN1 and ScXRN2/RAT1 (33,34). Orthologs of XRN1 and XRN2 have now been identified in many organisms including Arabidopsis (35). The Arabidopsis genome lacks an XRN1 homolog but codes for three XRNs, AtXRN2, AtXRN3 and AtXRN4, which are structurally similar to ScXRN2/RAT1 (35). AtXRN4 which lost the bipartite NLS, found on other XRN2 homologs, during duplication functions as a cytoplasmic exoribonuclease (35). XRN1 and XRN2 orthologs are widely conserved in eukaryotes and share two highly conserved regions called CR1 and CR2 in their N-terminal segments (36). Recently, a crystal structure of ScXRN1 in association with the 80S ribosome provided novel insights into the CTRD mechanism (37). Binding to the ribosome leads to a large rearrangement of ScXRN1 structure with three flexible loops in the exoribonuclease interacting directly with the ribosome. The most prominent rearrangement takes place in loop L3 located at the end part of the CR1 domain (37). This loop interacts with the mRNA emerging from the ribosome and is essential for the binding of ScXRN1 to ribosomes but is not essential for its catalytic activity. This loop was demonstrated to be conserved in Arabidopsis and essential for CTRD regulation (20). However, how XRN1/XRN4 association with the ribosome is controlled remains unknown.

XRN exoribonuclease activity can be inhibited by the secondary metabolite, 3ʹ- phosphoadenosine 5ʹ-phosphate (PAP), an intermediate of the sulfate assimilation pathway and a chloroplast retrograde signal accumulating during oxidative stress in plants (38–41). Under unstressed conditions, PAP is enzymatically degraded by the FRY1/SAL1 phosphatase (HAL2 in yeast) to form 5’AMP and Pi. In Arabidopsis, FRY1 loss-of-function leads to a constitutive overaccumulation of PAP that induces XRNs inhibition (38,40,41). Through grafting experiments, it was also proposed that the FRY1-PAP retrograde pathway could play a role in long distance signaling from shoot to root to modulate XRN activities in root (41). However, the detailed mechanism of this inhibition and the consequence on 5’-3’ mRNA turnover are unclear precisely regarding CTRD. Since FRY1 is functionally conserved, this would appear to be a conserved signaling pathway controlling XRN activity.

In addition to XRN exoribonuclease activity, PAP was also proposed to control DXO1 activity (42). DXO proteins are involved in 5’-end quality control and possess a deNADding activity (43,44). These proteins also present strong deNADding activity on mRNAs with non-canonical NAD^+^ cap that consists of nicotinamide adenine dinucleotide (45). In Arabidopsis, DXO1 activity was recently characterized. AtDXO1 crystal structure reveals conversed catalytic site but also plant-specific features (42). Two key amino acids, E394 and D396, were demonstrated to be involved in DXO1 exoribonuclease activity. Interestingly, catalytic activity does not contribute to several phenotypes observed in a *dxo1* knockout mutant (42). These phenotypes are linked to DXO1 N-terminal region, which is involved in the interaction with an RNA guanosine-7 methyltransferase (RNMT1) for mRNA guanosine cap methylation (46). However, its contribution to CTRD has never been tested.

Here, we analysed the regulation of CTRD by PAP and DXO1. Using *fry1* mutant and exogenous PAP treatment, we demonstrated that PAP could affect XRN4 association with polysomes and impair CTRD activity. Using 5’Pseq approach, we demonstrated that CTRD activity is more affected in *fry1* mutant than in *xrn4* mutant suggesting the involvement of other ribonucleases in the CTRD process. Finally, we demonstrated that in addition to XRN4, DXO1 is also involved in CTRD targeting NAD^+^-capped polysomal mRNAs.

## METHODS

### Growth conditions

All Arabidopsis lines have the Col-0 background. The *xrn4-5* T-DNA line was originally ordered from the Nottingham Arabidopsis stock centre (http://arabidopsis.info; SAIL_681_E01) and has been used in previous studies (21,31,32). The *fry1-4* was previously characterized in a mutagenesis screen (38). This mutant has a single point mutation at position 203 that generates a stop codon. This line was provided by H. Vaucheret (IJPB, Versailles, France). The DXO1(E394A/D396A) line was produced in (42). This transgenic line is expressing DXO1(E394A/D396A) fused to GFP in the *dxo1-2* (SALK_032903) background. This line was provided by J. Kufel (University of Warsaw, Warsaw, Poland). Seeds were sown on a 245x245mm square plate in a single row. Seedlings were grown vertically on synthetic Murashige and Skoog medium (Duchefa) containing 1% Sucrose and 0.8% plant agar at 22°C under a 16-h-light/8-h-dark regime. All the experiments were performed on 15-d-old seedlings. Roots and shoots were separated using a scissor at the basis of the hypocotyl and rapidly transferred to liquid nitrogen prior to RNA extraction.

### Total RNA extraction and 5’Pseq library preparation

Total RNA was isolated using Monarch Total RNA Miniprep Kit (New England Biolabs) according to manufacturer’s instructions. 5’Pseq libraries were prepared as described previously using 5 μg of total RNA as starting material (10). After sequencing, Raw reads for Read 1 were used and trimmed to 50 pb before mapping. Metagene analysis was performed using FIVEPSEQ software with standard parameters (8). The Terminational Stalling Index (TSI) was defined as the ratio of the number of 5’P read ends at the ribosome boundary (16–17 nt upstream from stop codons) to the mean number of 5’P read ends within the flanking 100 nt (20). Transcripts with a TSI value higher than 3 in Col0 were used to assess CTRD activity in the different mutant backgrounds. Two distinct batches of samples were prepared. Batch 1 includes Col0, *xrn4.5* and *fry1.4* shoot and root in three independent biological replicates (18 samples, PRJNA1185437). Batch 2 includes Col0, *xrn4.5* and DXO1(E394A/D396A) in three biological replicates (18 samples, PRJNA1189285). Each batch was sequenced independently. To compare NAD^+^-capped RNAs and DXO1 targets, supplemental dataset S4 from (47) was used. Only transcripts identified in both datasets were kept for the analysis. To test the significance of TSI distribution, a Wilcoxson-test was performed.

### Polysome profile and western-blot analysis

Polysome profiles and western-blot analysis were performed as described previously (21). Briefly, 400 mg of tissue power was homogenized in 1.2 ml of lysis buffer. After incubation 10 minutes on ice, lysate was clarified by centrifugation (10 minutes, 4°C, 16 000 g). 900 μl of crude extract was then loaded on a 15-60% sucrose gradient. After fractionation, proteins from fractions corresponding to polysomes were precipitated by adding 2 volumes of absolute ethanol. After 6 hours at 4°C and centrifugation, protein pellets were washed 5 times with ethanol 70%. Finally, pellets were resuspended in Laemmli 4X buffer. XRN4 antibody (31) was used at 1/1000th dilution. RPL13 antibody was purchased (Agrisera) and used at 1/5000th dilution. Primary antibody was incubated overnight at 4°C under constant agitation. A horseradish peroxidase-coupled antibody was used as secondary antibody. Signal was revealed with the Immobilon-P kit (Millipore).

### RT-ddPCR after transcription inhibition

*In vivo* transcription inhibition was performed as in (20,48). Plantlets were transferred horizontally in an incubation buffer (15 mM sucrose, 1 mM Pipes pH 6.25, 1 mM KCl, 1 mM sodium citrate, 1 mM cordycepin). For PAP treatment, 1 mM ATP and 1 mM PAP were added in the incubation buffer as described (49). The time-course experiment was performed 2 hours after daybreak. Roots and shoots were collected separately at 0, 30, 60 and 120 min after transcription arrest and rapidly transferred to liquid nitrogen prior to RNA extraction. 500 ng of total RNA was reverse-transcribed using SuperScript IV kit and random primers (Thermo Scientific). cDNAs were then diluted 50-fold prior to ddPCR analysis. ddPCR was performed as described previously (50). Primers used for ddPCR are listed on Supplemental Table 1. mRNA half-life determination was performed with three independent biological replicates.

## RESULTS

### FRY1 inactivation affects co-translational mRNA decay

XRN exoribonuclease activity can be inhibited by 3’-phosphoadenosine 5’-phosphate (PAP) (40). Under unstressed conditions, PAP is rapidly degraded by the FRY1 phosphatase. FRY1 loss-of-function leads to constitutive PAP accumulation and this accumulation inhibits XRN activity (38,40,41). However, the detailed mechanism of this inhibition and the consequences for mRNA turnover are unclear. Here, we tested if FRY1 through PAP accumulation could be involved in CTRD modulation. First, we investigated XRN4 accumulation in polysomes (as a read-out of CTRD activity) in shoot and root of 15-d-old Col0 and *fry1* mutant seedlings (Figure 1). After polysome profiling (Figure 1A-B), fractions corresponding to polysomes were collected and analysed via western blotting with an XRN4-specific antibody (Figure 1C-D). While the XRN4 signal is similar in each input fraction, its accumulation in polysomes seems to be affected in both *fry1* shoot and root, suggesting that FRY1 inactivation can impair CTRD activity. To test this hypothesis, we performed a 5’Pseq approach in Col0, *xrn4* and *fry1* mutants to assess CTRD activity (Figure 2 and Supplemental Figure 1). 5’Pseq was performed on 3 biological samples, revealing reproducible results between genotypes and organs (Supplemental Figure 1). To assess CTRD activity, we first performed a metagene analysis around stop codons in each condition (Figure 2A). A clear overaccumulation of 5’P reads at position -16/-17nt before stop codons is observed in Col0 shoot and root, a hallmark of active CTRD. This peak drastically decreased in both *xrn4* and *fry1* mutants (Figure 2A). As metagene analysis can biases the signal to higher abundance transcripts, we determined CTRD signal at the individual transcript level using the Terminational Stalling Index (TSI, (19,20,22)). The higher the TSI, the greater the CTRD activity. This index was determined in shoot and root respectively for Col0, *xrn4* and *fry1* mutants (Figure 2B-C). Only transcripts with a TSI value higher than 3 in Col0 were considered as CTRD targets (20,22). In *xrn4* and *fry1*, the TSI distribution drastically decreases in both shoot and root. Interestingly, this decrease is significantly stronger in *fry1* than in *xrn4* in both organs. We then compared XRN4 and FRY1 CTRD targets. To do so, we considered transcripts that present lower TSI values in *fry1* and/or *xrn4* compared to Col0 as FRY1 and/or XRN4 CTRD targets (Figure 2B-C). Strong overlap between XRN4 and FRY1 CTRD targets was observed in both organs with respectively 5 571 (89 % and 91% of FRY1 and XRN4 targets respectively) and 4 403 (85.6 % and 86.7 % of FRY1 and XRN4 targets respectively) common targets in shoot and root (Figure 2B-C, Supplemental Tables 2 and 3). These data suggest that FRY1 inactivation, likely through PAP accumulation can inhibit XRN4-CTRD activity. As CTRD is more repressed in *fry1* than in *xrn4*, these data also suggest that other enzymes may be involved in the CTRD pathway.

**Figure 1 :**
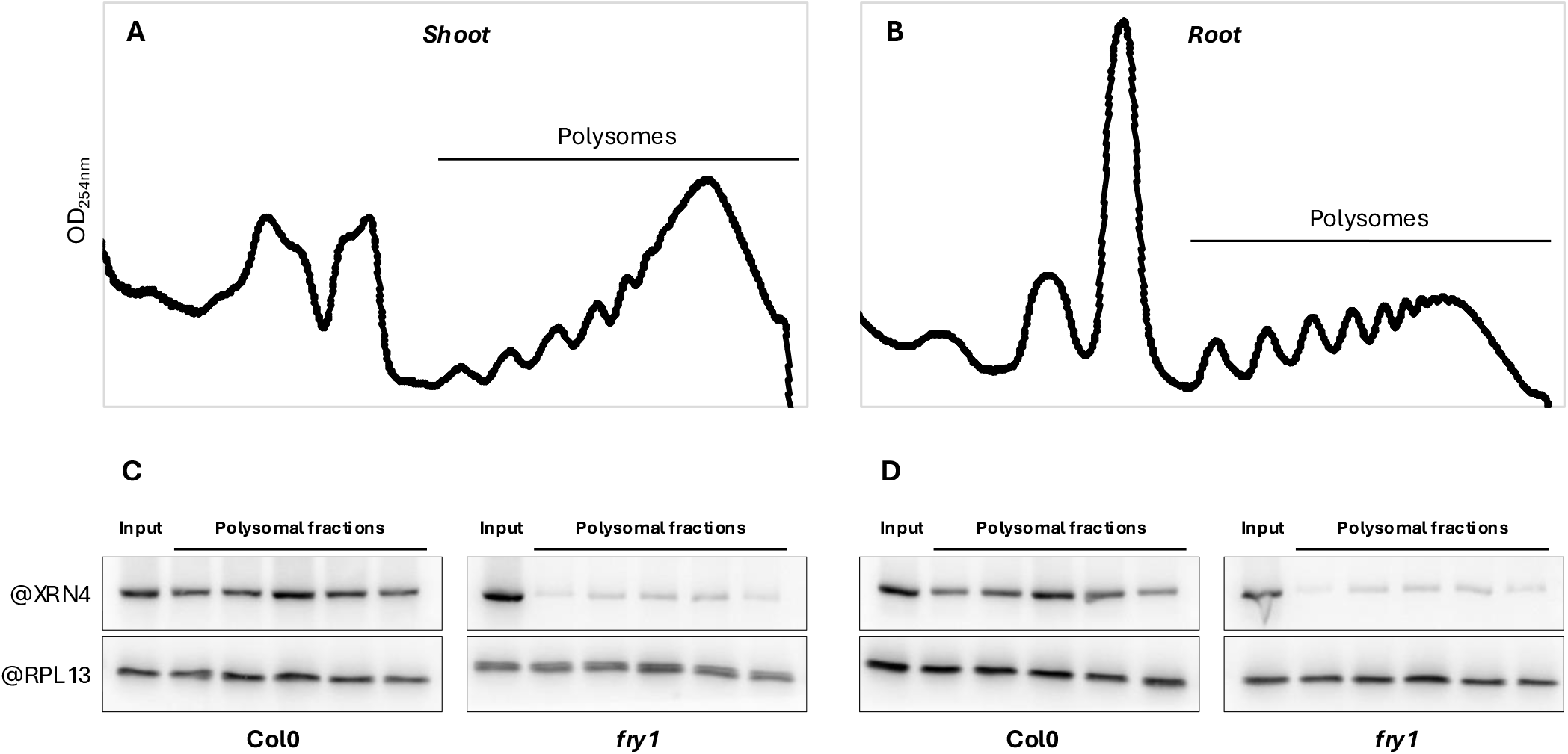
XRN4 accumulation to polysomes is impaired in *fry1* mutant. Polysomal extracts prepared from Col0 or *fry1* shoot (A) and root (B) were fractionated on a sucrose gradient. C-D. Total proteins extracted from polysomal and input fractions were analysed by immunoblotting using XRN4 or RPL13 specific antibodies. The four blots were prepared and analysed simultaneously. The same quantities of tissues were used for each condition (e.g. 300 mg of biomass).

**Figure 2 :**
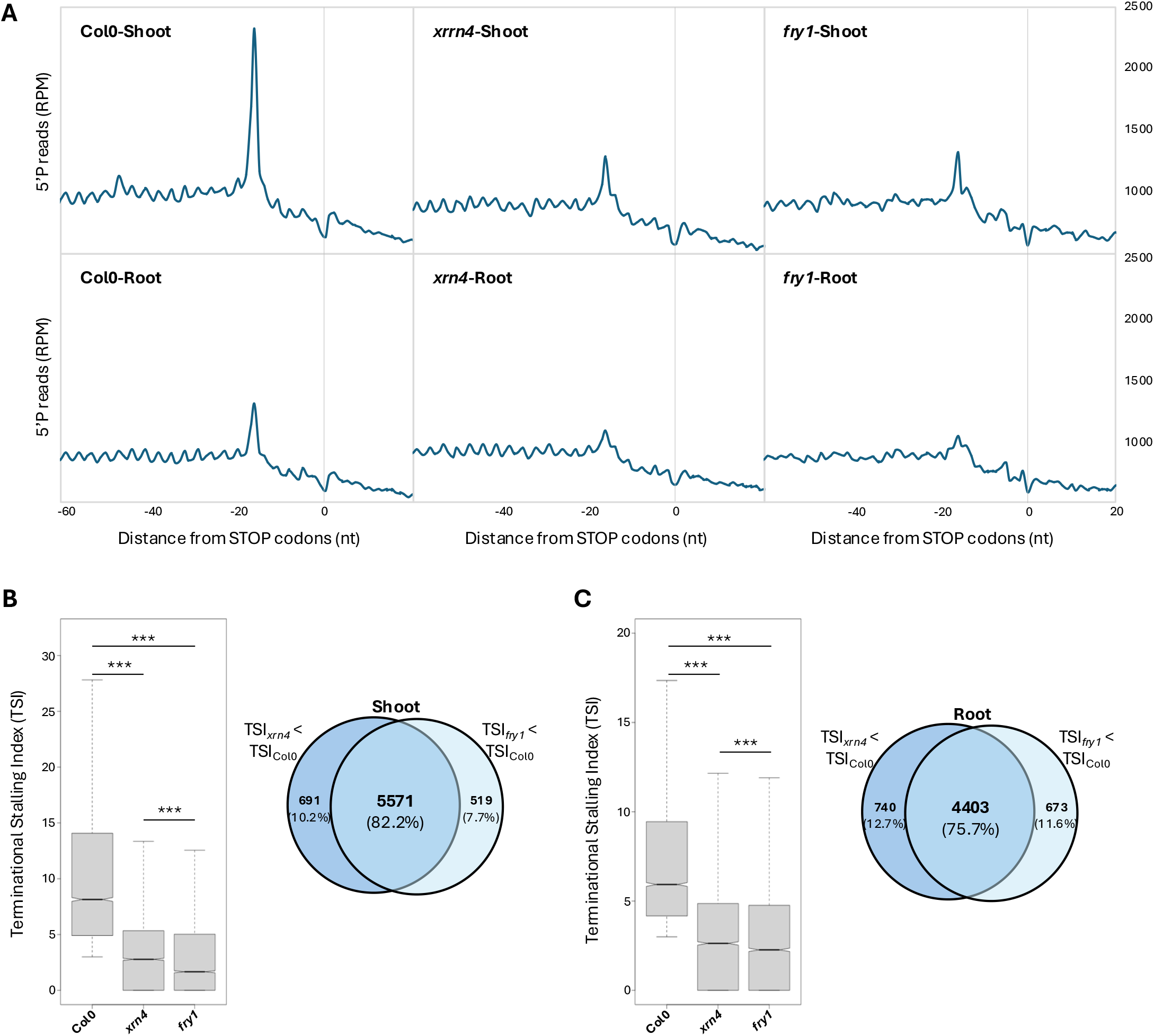
FRY1 inactivation affects co-translational mRNA decay in both shoot and root. A. Metagene analysis of 5′P reads accumulation around stop codons. The different profiles are representative of 3 biological replicates. B. Distribution of Terminational Stalling Index (TSI) in Col0, *xrn4* and *fry1* and comparison of XRN4 and FRY1 targets in shoot. C. Distribution of Terminational Stalling Index (TSI) in Col0, *xrn4* and *fry1* and comparison of XRN4 and FRY1 targets in roots. A transcript was considered as a target when the TSI value in *xrn4* and/or *fry1* is lower than in Col0. N = 3 biological replicates.

### PAP treatment affects XRN4 accumulation to polysomes and mRNA stability

As a constitutive accumulation of PAP is observed in *fry1* mutant (39), we tested whether exogenous PAP treatment can also affect CTRD activity. To do so, whole seedlings were treated *in vivo* with 1 mM exogenous PAP. Exogenous ATP was also added as is a known co-substrate for the PAP transporter (49,51). After 1 hour of treatment, shoots and roots were collected separately. After polysome fractionation, XRN4 accumulation in polysomes was tested via western blotting (Figure 3A). Interestingly, exogenous PAP treatment reduces XRN4 accumulation in polysomes in both organs without affecting total XRN4 signal. This result suggests that PAP can affect CTRD activity. To test this hypothesis, we measured the mRNA half-lives of candidate CTRD targets after cordycepin treatment supplemented with 1 mM PAP (Figure 3B-C). We choose two transcripts targeted by CTRD that present lower TSI in *fry1* and *xrn4* mutants. For these two candidate targets, PAP treatment significantly affects mRNA stability in both organs. For At2g21350 transcript, mRNA half-life varied from 20.8 minutes to 141.5 minutes and from 24.1 minutes to 188.6 minutes after PAP treatment in shoot and root respectively. At1g66300 transcript followed the same trend with an mRNA half-life that varies from 17.2 minutes to 58.1 minutes and from 25.7 minutes to 58.8 minutes after PAP treatment in shoot and root respectively (Figure 3B-C). Collectively, these data suggest that PAP can regulate CTRD activity in shoot and root.

**Figure 3 :**
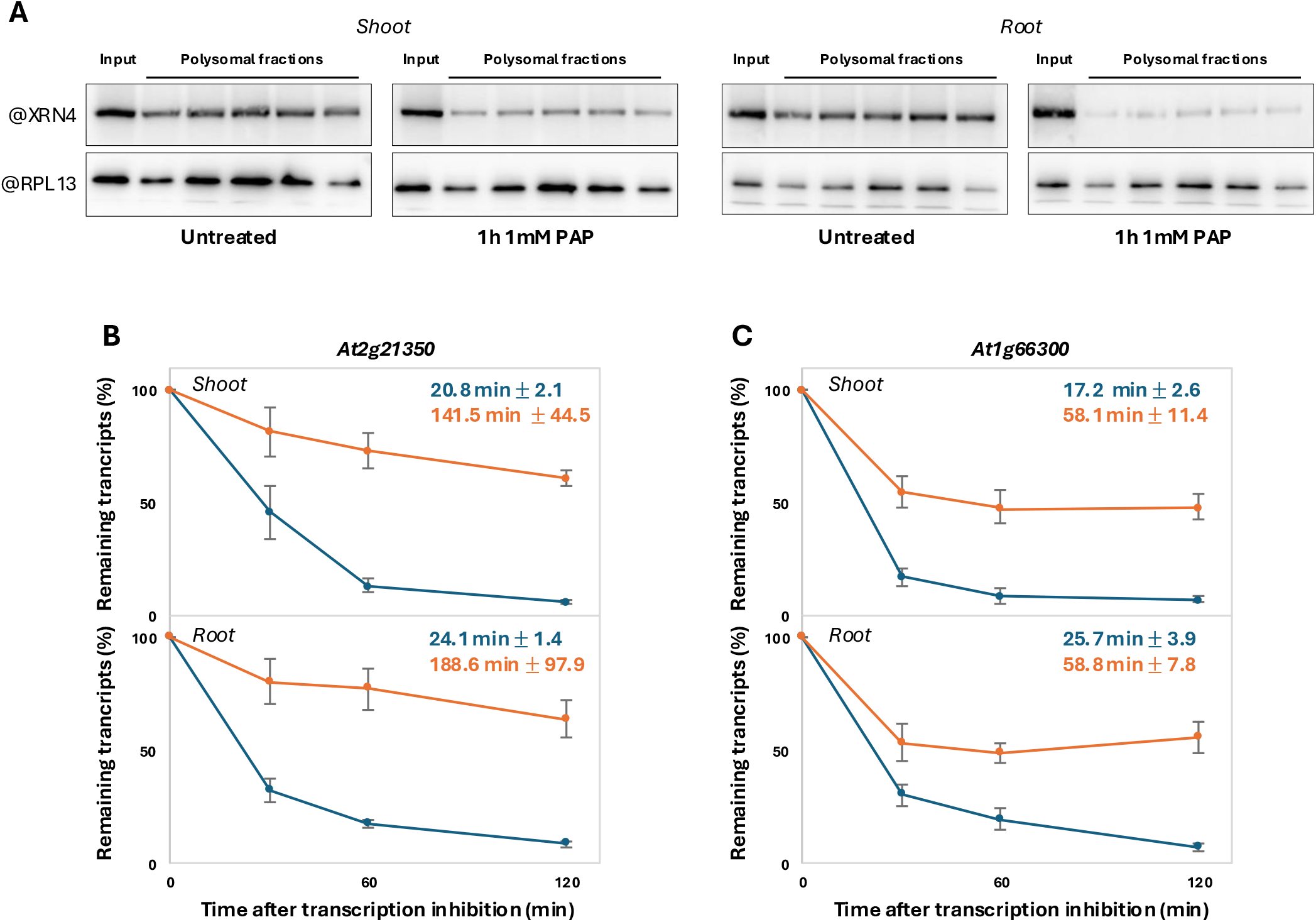
PAP treatment affects XRN4 accumulation to polysomes and mRNA stability. A. Total proteins extracted from polysomal and input fractions prepared from (A) Col0 shoot or (B) Col0 root incubated for 1 hour on liquid MS medium (untreated) or 1 hour on liquid MS medium supplemented with 1 mM PAP. The four blots were prepared and analysed simultaneously. The same quantities of tissues were used for each condition (e.g. 300 mg of biomass). B, C. mRNA stability was determined *in vivo* after 1 mM cordycepin treatment (blue line) or after 1 mM cordycepin and 1 mM PAP treatment (orange line) followed by RT-ddPCR. Half-lives are indicated on each graph. N = 3 biological replicates.

### DXO1 catalytic inactivation affects co-translational mRNA decay

Recently, it was reported that in addition to XRN activity, PAP can also inhibit DXO1 exoribonuclease activity (42). DXO1 is implicated in the removal of non-canonical NAD^+^ cap and in RNA turnover (42,52). However, its contribution to CTRD has never been tested. As the *dxo1* knockout mutant presents drastic phenotypes not directly linked to its catalytic activity (42,46), we used a catalytically inactive DXO1, DXO1(E394A/D396A) expressed in a *dxo1-2* knockout mutant to test its role in CTRD (42). In comparison to Col0 and *xrn4*, we performed a 5’Pseq analysis in DXO1(E394A/D396A) shoot and root (Figure 4). 5’Pseq was performed on 3 biological samples, revealing reproducible results between genotypes and organs (Supplemental Figure 2). Metagene analysis around stop codons reveals a decrease of the peak at position -16/-17nt before stop codons in DXO1(E394A/D396A) line compared to Col0 but to a lesser extent than in *xrn4* in both shoot and root (Figure 4A). To confirm this moderate effect, we analysed the TSI distribution in each genotype (Figure 4B-C). In both organs, the TSI distribution significantly decreased in DXO1(E394A/D396A) and *xrn4* lines. Interestingly, the decrease is significantly lower in *xrn4* than in DXO1(E394A/D396A) line in shoot and in root. However, a clear overlap between the DXO1 and XRN4 CTRD targets was observed with 92.8.% and 87.6% of DXO1 targets are also XRN4 targets in shoot and root respectively (Figure 4B-C, Supplemental Tables 4 and 5). These data suggest that XRN4 can potentially also degrade DXO1 targets after a NAD^+^ cap removal.

**Figure 4:**
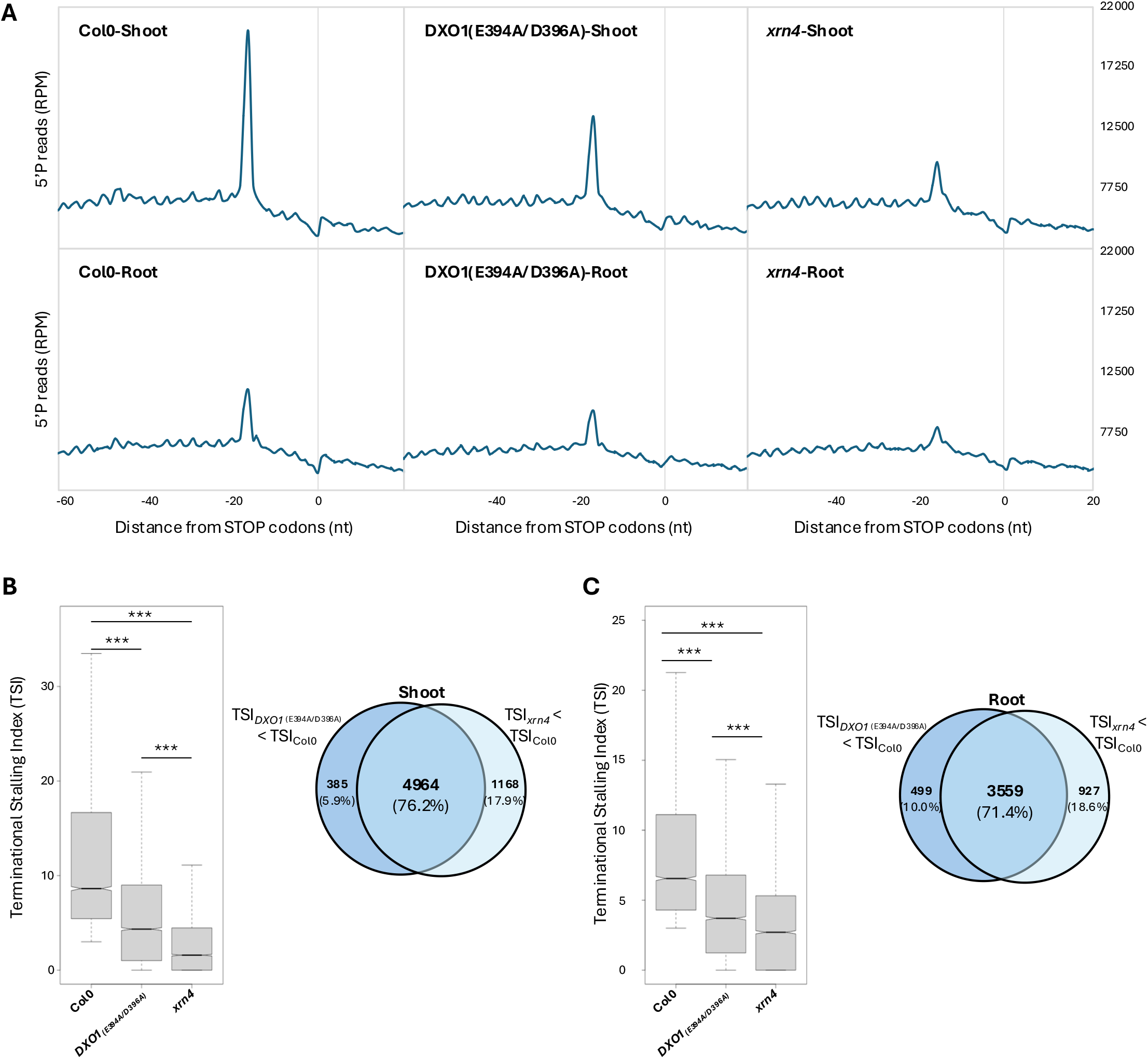
DXO1 catalytic inactivation affects co-translational mRNA decay in both shoot and root. A. Metagene analysis of 5′P reads accumulation around stop codons. The different profiles are representative of 3 biological replicates. B. Distribution of Terminational Stalling Index (TSI) in Col0, DXO1(E394A/D396A) and *xrn4* and comparison of XRN4 and DXO1 targets in shoot. C. Distribution of Terminational Stalling Index (TSI) in in Col0, DXO1(E394A/D396A) and *xrn4* and comparison of XRN4 and DXO1 targets in root. A transcript was considered as a target when the TSI value in DXO1(E394A/D396A) and/or *xrn4* is lower than in Col0. N = 3 biological replicates.

### DXO1 co-translational mRNA decay targets are identified as NAD^+^-capped RNAs in polysomes and targeted by FRY1

As DXO1 is known to be involved in the removal of non-canonical NAD^+^ cap, we tested whether DXO1-identified co-translational mRNA decay targets can harbour a NAD^+^ cap. Recently, a survey of NAD^+^-capped RNAs was performed using NAD captureSeq approach (47). The authors reported that NAD^+^-capped mRNAs can be identified in polysomes. Thus, we compared NAD^+^-capped RNAs identified in polysomes with DXO1 co-translational mRNA decay targets. Given that the NAD captureSeq approach was performed on seedlings, we compared the data with DXO1 CTRD targets in shoot. Moreover, we retained only the transcripts identified in both datasets resulting in a total of 3 637 transcripts harbouring a NAD^+^ cap in polysomes and 6 321 DXO1 targets (TSI_DXO1(E394A/D396A)_ < TSI_Col0_). Interestingly, 71.1% (2 586/3 637, p-value 2.6^e-95^) of the NAD^+^-capped polysomal mRNAs were identified as DXO1 CTRD targets in our dataset (Figure 5A). In order to determine biological processes targeted by DXO1 co-translational mRNA decay, a Gene Ontology (GO) analysis was performed on NAD^+^ specific polysomal RNA DXO1 CTRD targets (N=2 586, Figure 5B). Among the different GO terms identified, many terms associated with “Response to stimulus” were enriched in the dataset (GO=00508896, 4.22^e-37^, Figure 5B). These terms can be classified into two subterms, “Response to chemical” (GO:0042221, 9.94^e-30^) and “Response to abiotic stimulus” (GO:0009628, 4.71^e-40^). Among them, the GO term “Response to abscisic acid” was significantly enriched (GO:0009737, 2.78^e-06^). Interestingly, DXO1 has been reported to regulate mRNA stability of ABA-related NAD^+^-capped mRNAs (53). As DXO1 is also sensitive to PAP, we compared DXO1 targets to FRY1 targets in shoot and root (Figure 5C-E). To do so, we selected transcripts with a lower TSI value in DXO1(E394A/D396A) and/or *fry1* lines than in Col0 in both organs. Venn diagram analysis revealed strong overlap between the DXO1 and FRY1 targets with 85.9% of DXO1 targets are also FRY1 targets. Additionally, the TSI distribution in both organs reveals a stronger effect in *fry1* than in DXO1(E394A/D396A) (Figure 5D-E). Taken together, these data suggest that DXO1 can target NAD^+^-capped mRNAs to CTRD, which can also be targeted by other exoribonucleases after NAD^+^ cap removal.

**Figure 5 :**
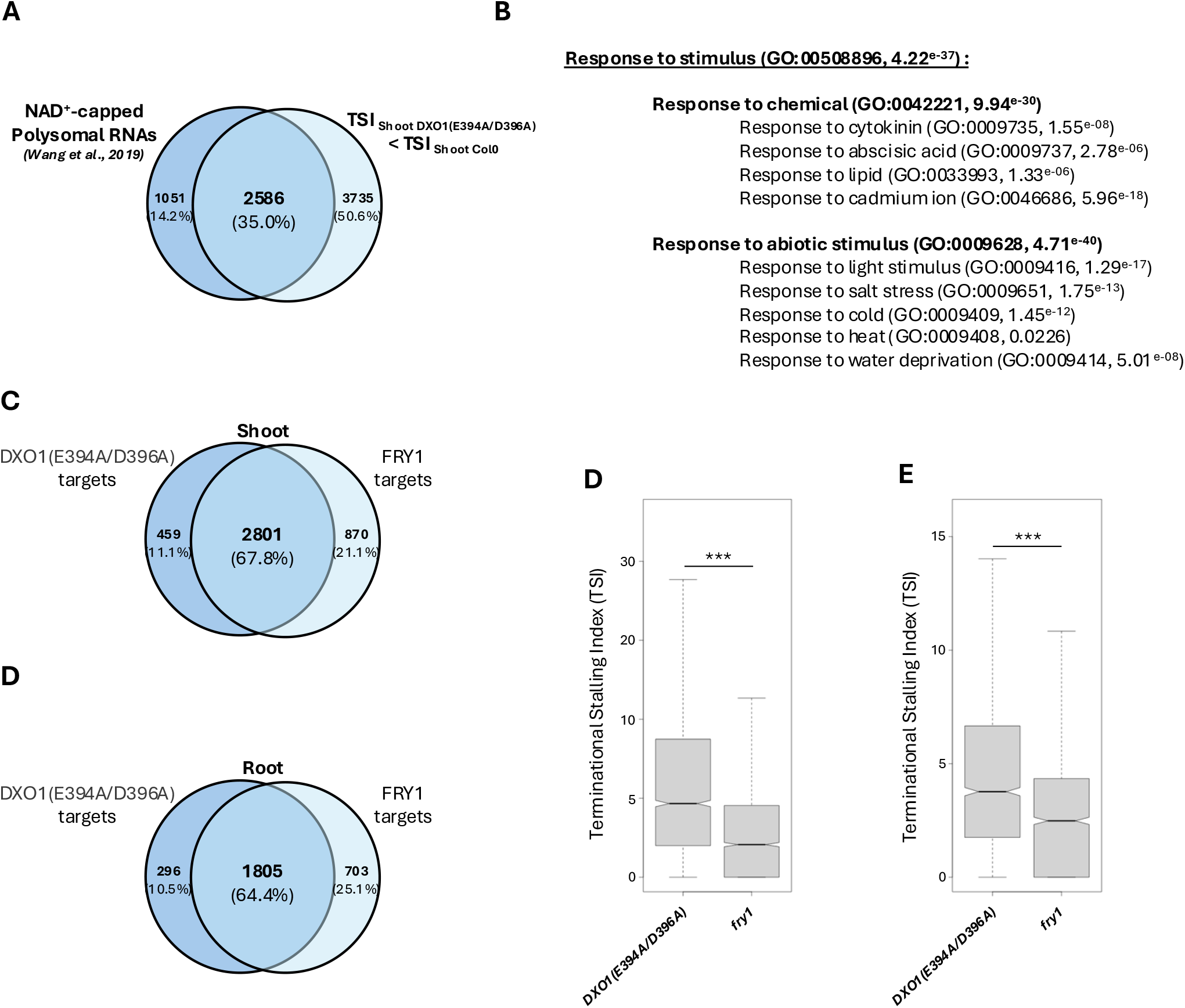
DXO1 co-translational mRNA decay targets are identified as NAD^+^-capped RNAs in polysomes and targeted by FRY1. A. Venn diagram representation of the number of NAD^+^-capped transcripts identified in polysomes according to (47) and DXO1 co- translational mRNA decay targets identified in shoot. Transcripts with a TSI value higher than 0 in Col0 were kept for the analysis. To test the significance of the overlap, hypergeometric test was performed (p-value 2.6^e-95^). B. List of response to stimulus GO terms that are enriched in DXO1 co-translational mRNA decay targets (N = 2 586). C, D. Comparison of DXO1(E394A/D396A) and FRY1 targets in shoot and root respectively (A transcript was considered as a target when the TSI value in DXO1(E394A/D396A) and/or *fry1* is lower than in Col0). D, E. Distribution of the TSI for the common targets of DXO1(E394A/D396A) and FRY1 in shoot and root respectively (N = 2 081 in shoot and N = 1 805 in root).

## DISCUSSION

Since its discovery, the CTRD pathway has been found to globally shape the whole transcriptome in several organisms (11,20,22,24,26,54). However, details about its regulation are still rare in the literature. Here, using 5’Pseq approaches and mRNA half-life determination, we reported that PAP and DXO1 are involved in CTRD regulation in both shoot and root in Arabidopsis.

XRN4 was described as the main enzyme involved in CTRD in Arabidopsis (20,21,23,24). However, we and others have reported that a slight CTRD activity is still present in the *xrn4* mutant (20,21,23). In fact, the 5’P reads accumulation 16/17 nucleotide before stop codons is not totally abolished in the *xrn4* mutant (20,21). Moreover a 3-nt periodicity is still observed in the *xrn4* mutant suggesting that additional pathways can contribute to this periodicity. Recent studies have proposed that ribosome dynamics can also be linked with endoribonuclease activity. For example, in yeast, ribosome collisions can trigger mRNA cleavage at ribosome boundaries via the endoribonucleases Cue2 and SmrB (55,56). In Arabidopsis, only one endoribonuclease was recently characterized (57,58). However, its link with ribosome dynamics has not been assessed. CTRD mediated by other exoribonucleases could also explain the remaining slight CTRD activity in the *xrn4* mutant. Arabidopsis genome encodes three XRN genes, XRN2, XRN3 and XRN4. Recently, a large overview of the Arabidopsis mRNA degradome was obtained via degradome sequencing of the different *xrn* mutants in addition to *fry1* (23). It appears that XRN2 and XRN3 have no redundant functions with XRN4 in the CTRD. This analysis was performed only on single mutants and cannot reveal compensate mechanism involving XRN2 and/or XRN3. Additionally, this analysis revealed that mRNA decay is more repressed in *fry1* than in *xrn4* (23). These data are consistent with our findings. In fact, through TSI analysis, our data revealed that XRN4 and FRY1 share similar targets with a stronger effect in *fry1* than in *xrn4* (Figure 2).

To test the potential involvement of other exoribonucleases in CTRD, we tested the contribution of DXO1 to this process. In Arabidopsis, DXO1 was demonstrated to possess deNADding and exoribonuclease activities (42,59). DXO1 inactivation induces pleiotropic phenotypes such as growth defects, pale green coloration or insensitivity to ABA (42,53,59). Our data revealed that DXO1 contributes to CTRD but to a lesser extent than XRN4 does (Figure 4). Using published NAD captureSep data, we also demonstrated that DXO1 targets possess NAD^+^ cap. These data suggest that XRN4 can also target NAD^+^-capped transcripts after a deNADding process. Interestingly, DXO1 is a distributive enzyme while XRN4 harbours processive activity (60). It is tempting to propose that after deNADding, XRN4 can compete with DXO1 for co-translational mRNA decay. Recently in Arabidopsis, the absence of XRN4- CTRD pathway in roots was proposed to result in a large feedback mechanism by an unknown decay pathway (20) suggesting that other exoribonucleases can also compensate for the absence of XRN4-CTRD. Finally, the moderate impact of DXO1 on global CTRD can be explained by its selectivity for only NAD^+^-capped mRNAs.

Our data also reveal that PAP can affect CTRD activity. A biochemical characterization of Human XRN1 in interaction with PAP was recently performed (61). The crystal structure reveals that PAP bounds to the active site of XRN1, the adenine base of PAP forms a π-stacking interaction with the amino acid His41 (61). Interestingly, this amino acid appears conserved and is also important for XRN1 interaction to the ribosome in yeast and drosophila (37) suggesting that PAP can interfere the interaction XRN1/Ribosome and impaired CTRD activity. This amino acid is also present in AtXRN4 (His61) suggesting a conserved regulatory mechanism (20).

The GO analysis also revealed that DXO1 CTRD targets are enriched in mRNAs responsible for chemical responses, particularly mRNAs linked to the ABA response (Figure 5B). This term is consistent with the proposed role of DXO1 in ABA-related transcript stability (53). In addition, GO terms linked to stress response appears also enriched. Future experiments will be needed to determine the direct link between DXO1-CTRD pathway and stress response in Arabidopsis.

## CONCLUSIONS

In conclusion, our study revealed that the CTRD pathway can be modulated by PAP and triggered by DXO1 in Arabidopsis. For the first time, we demonstrated the role of another exoribonuclease, DXO1, in the regulation of this pathway, probably by targeting NAD^+^-capped mRNAs. Finally, this study paves the way for studying the regulatory importance of CTRD in plants.

## LIST OF ABBREVIATIONS

CTRD: Co-Translational mRNA Decay
DXO1: Decapping and exoribonuclease protein 1
FRY1: FIERY1
NAD: Nicotinamide Adenine Dinucleotide
PAP: 3ʹ-PhosphoAdenosine 5ʹ-Phosphate
XRN4: Exoribonuclease 4

## DECLARATIONS

### Ethics approval and consent to participate

Not Applicable

### Consent for publication

Not Applicable

### Availability of data and materials

The 5’Pseq datasets generated in this study have been deposited in the Short Read Archive with the accession code PRJNA1185437 and PRJNA1189285.

### Authors’ contributions

Rémy Merret conceived and designed the project. Rémy Merret and Adrien Cadoudal performed the experiments. Marie-Christine Carpentier, Anne-Elodie Receveur and Rémy Merret analyzed the data. Rémy Merret wrote the manuscript. All authors read and approved the manuscript.

## Acknowledgements

This study is set within the framework of the “Laboratoires d’Excellences (LABEX)” TULIP (ANR- 10-LABX-41) and of the “École Universitaire de Recherche (EUR)” TULIP-GS (ANR-18-EURE- 0019). We would like to thank Joanna Kufel and Monika Zakrzewska-Placzek for DXO1(E394A/D396A) line and Hervé Vaucheret for *fry1.4* line.

---

**Supplemental Table 1.** List of primers using for RT-ddPCR.

**Supplemental Table 2.** List of genes identified as co-translational mRNA decay targets in shoot for Col0, *xrn4* and *fry1*.

**Supplemental Table 3.** List of genes identified as co-translational mRNA decay targets in root for Col0, *xrn4* and *fry1*.

**Supplemental Table 4.** List of genes identified as co-translational mRNA decay targets in Shoot for Col0, *DXO1(E394A/D396A)* and *xrn4*.

**Supplemental Table 5.** List of genes identified as co-translational decay mRNA targets in root for Col0, *DXO1(E394A/D396A)* and *xrn4*.

**Supplemental Figure 1 :**
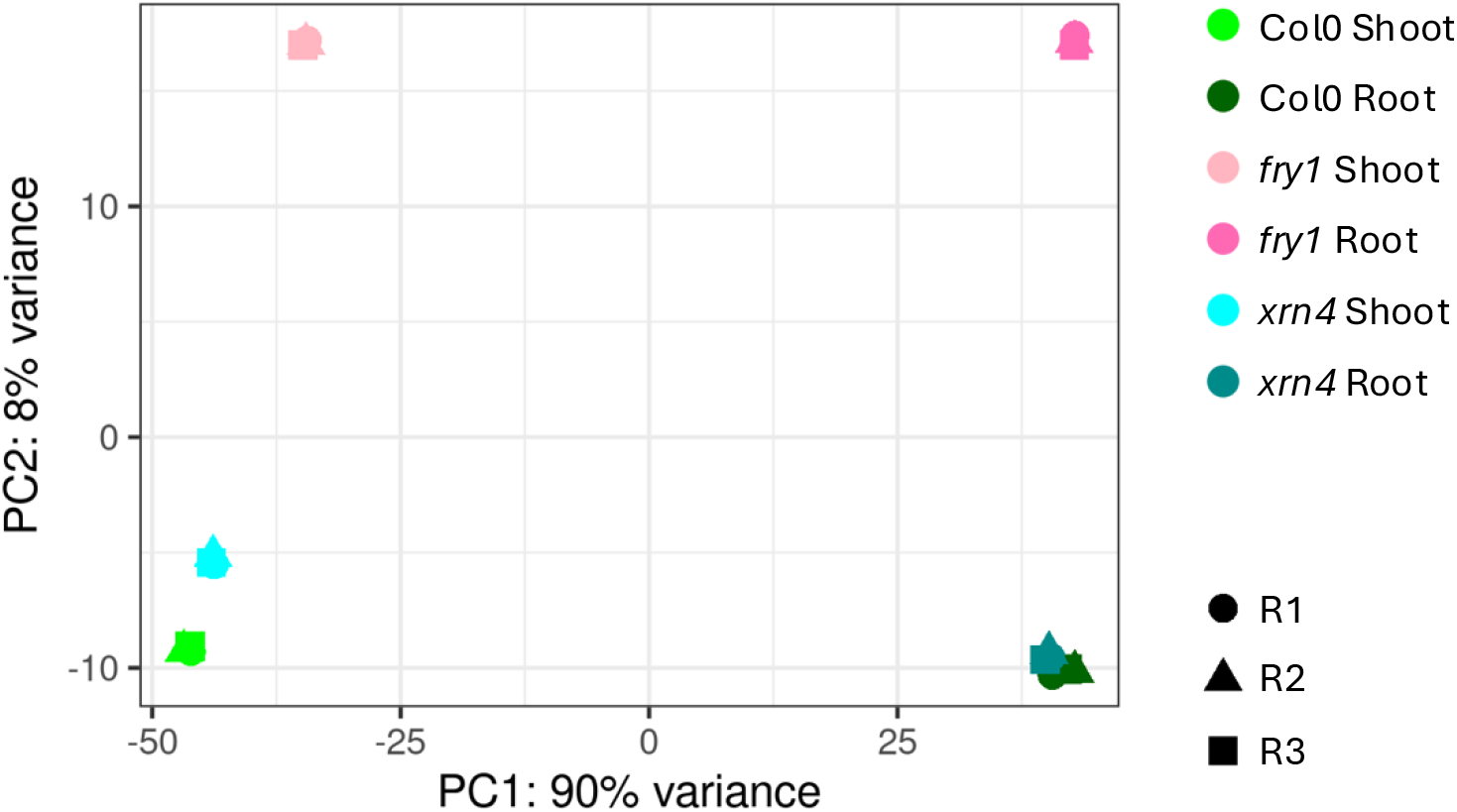
Principal component analysis for Col0, *xrn4* and *fry1* 5’Pseq samples. N = 3 biological replicates per genotype per organ.

**Supplemental Figure 2 :**
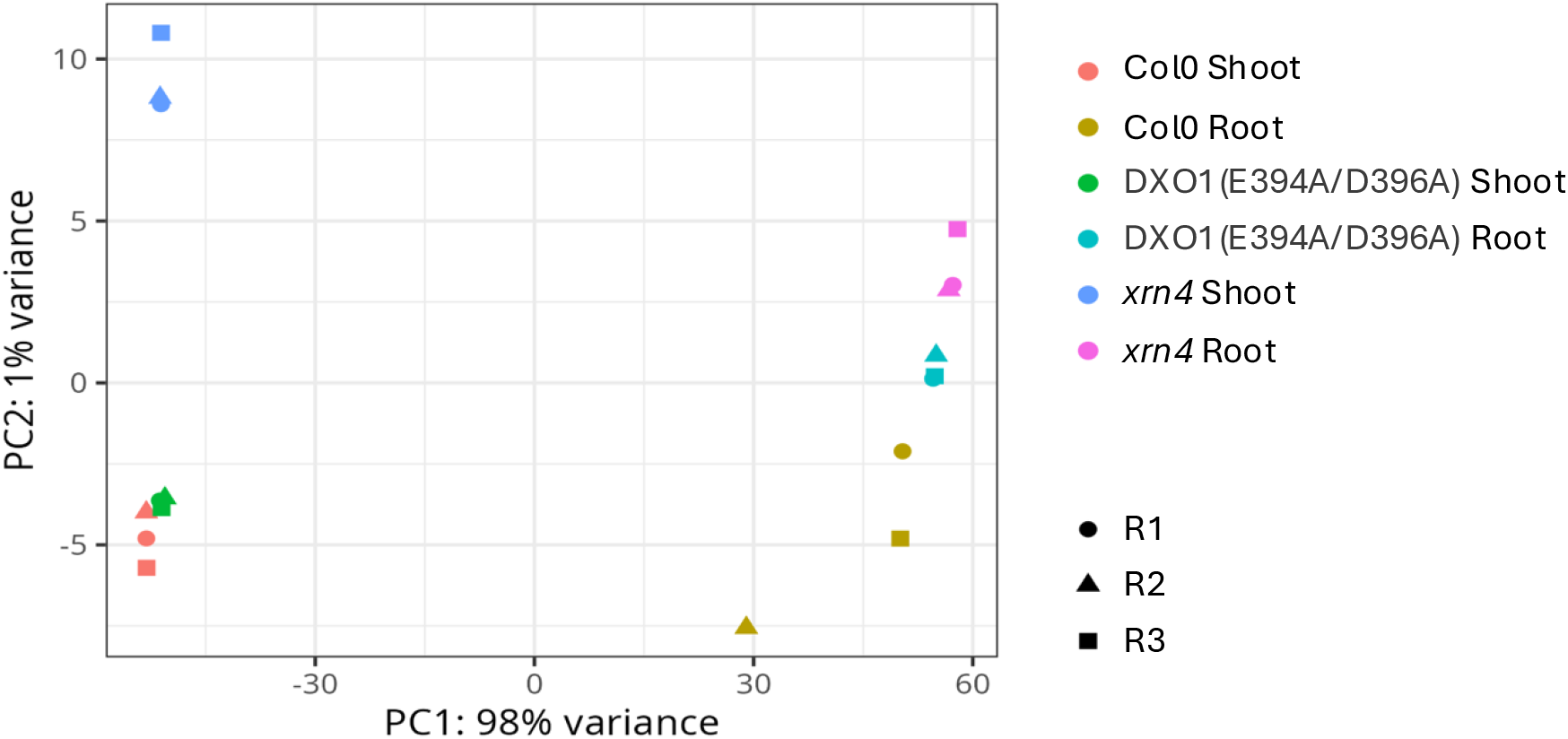
Principal component analysis for Col0, *xrn4* and DXO1(E394A/D396A) 5’Pseq samples. N = 3 biological replicates per genotype per organ.

